# An astro-neural-field model with application to cortical spreading depolarization

**DOI:** 10.64898/2026.06.10.731347

**Authors:** E. Baspinar, D. Avitabile, C. Nouveau, M. Desroches, F. Campillo, M. Mantegazza

**Affiliations:** MathNeuro Team, Inria Branch of the University of Montpellier, Montpellier, France; Department of Mathematics, Vrije Universiteit Amsterdam, Amsterdam, The Netherlands; Amsterdam Neuroscience - System and Network Neuroscience, Amsterdam, The Netherlands; Université Côte d’Azur, Valbonne-Sophia Antipolis, France; CNRS UMR7275, Institute of Molecular and Cellular Pharmacology (IPMC), Valbonne-Sophia Antipolis, France; Inserm U1323, Valbonne-Sophia Antipolis, France

**Keywords:** Neural field, Astrocyte, Migraine, Spreading depolarization

## Abstract

We present a novel astro-neural-field population model with application to migraine-related cortical spreading depolarization. The model is composed of four spatio-temporal state variables: excitatory and inhibitory membrane potentials, astrocytic potassium uptake recruitment, and extracellular potassium concentration. Extending a previous neural field model, we incorporate activity-dependent astrocytic potassium clearance via a nonlinear term coupled to astrocyte dynamics. The astrocyte transfer function, like its neural counterpart, exhibits three regimes governed by extracellular potassium, capturing its effect on clearance. This yields a more comprehensive framework, better fits experimental data, and provides new insights into the mechanisms of cortical spreading depolarization.

## 1 Introduction

In the present work, we extend a previously proposed computational model of neural dynamics observed in a pathological phenomenon known as cortical spreading depolarization (CSD) (Baspinar et al. 2025). This extension generalizes CSD-related neural dynamics to astrocytic dynamics.

CSD is a pathological brain phenomenon observed in several neurological disorders. It is a wave of transient intense neural firing followed by neural depolarization, which begins focally and propagates slowly across the cortex by inducing several pathological conditions. This wave of intense neural firing is followed by a sustained neural depolarization and an increase of extracellular potassium (K^+^) concentration in the range of tens of mM. The depolarization induces a prolonged suppression of neural activity by blocking neural firing; see Fig. 1 for experimental recordings.

**Fig. 1.**
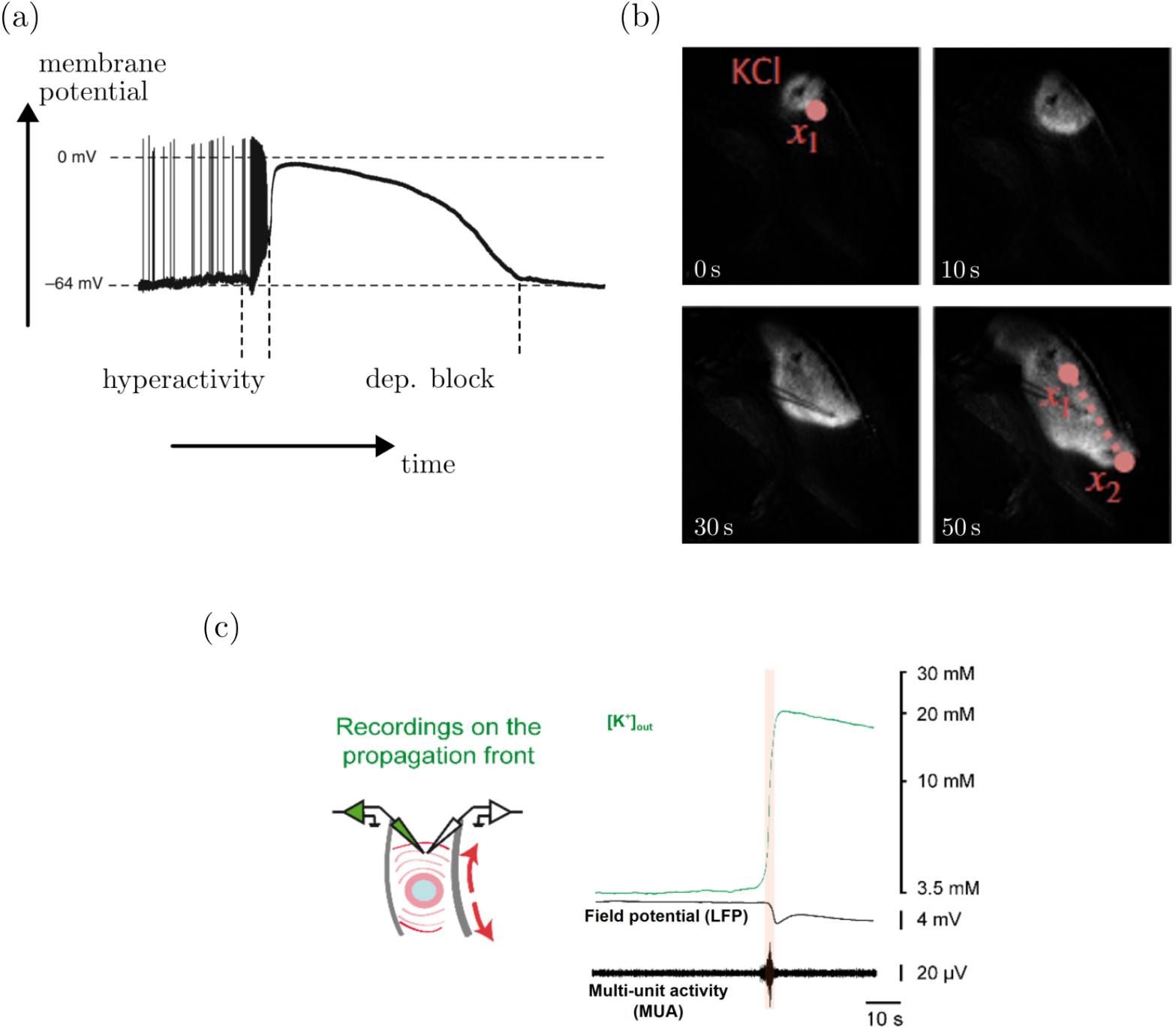
CSD initiation and propagation. (a) Intracellular electrode recording from a neuron experiencing CSD (Ghadiri et al. 2012). Intense spiking (hyperactivity) is followed by the depolarization block suppressing the spiking and maintaining the membrane potential between 0 and − 30 mV (see (Pietrobon and Moskowitz 2014) for a review.). (b) Four instants of CSD propagation recorded via optical imaging (Baspinar et al. 2025). Bright and dark pixels represent high and low neural activity regions, respectively. The region where potassium chloride (KCl) drops are applied is highlighted as KCl. Two electrodes are fixed at *x*_1_ and *x*_2_, which correspond to the initial and final positions of the propagating wavefront, respectively. The dashed line shows the propagation direction of the CSD wavefront. KCl ≈ 130 mM is within the lower range commonly used in the CSD field, where concentrations up to 1 M have been employed to ensure reliable initiation (Pietrobon and Moskowitz 2014). This concentration is not physiologically impossible: local cell lysis (e.g., after trauma) can transiently expose extracellular space to intracellular K^+^ concentrations (≈130 mM). Moreover, the propagating CSD wavefront that is studied here is determined by the intrinsic network and ionic dynamics and is not directly influenced by the initial KCl concentration at the induction site. c) Quantification of extracellular K^+^ concentration ([K^+^]_out_) dynamics with associated local field potential recordings and multi-unit activities (MUA) recorded on the CSD propagation front in cortical slices (from (Chever et al. 2021)). Left: Diagram illustrating the recordings. Right: Representative traces; the vertical orange area highlights the period of paroxysmal MUA, which corresponds to the transient intense neuronal firing that precedes the long lasting depolarization block in CSD. In these experiments extracellular potassium increased up to 40 mM. In the literature, concentrations up to 100 mM have been reported (Mantegazza et al. 2010), with average values in the range 30-60 mM (Pietrobon and Moskowitz 2014).

CSD was first observed in a rabbit brain (Leão 1944). Subsequently, it has been observed in other animal models (James et al. 1999). These studies showed that CSD has important roles in several neurological disorders, such as traumatic brain injury (Lauritzen et al. 2011; Soldozy et al. 2020), epilepsy (Dreier et al. 2011), and in particular migraine (Lauritzen 1994; Lauritzen et al. 2011; Major et al. 2020).

Despite the increasing clinical importance of CSD, it has only recently been detected in humans (Major et al. 2020; Mayevsky et al. 1996; Strong et al. 2002). Moreover, it remains difficult to detect it noninvasively in humans. This calls for mathematical models to investigate the biological mechanisms underlying CSD. A particularly important aspect is that computational models can be used to simulate experimental protocols, providing a powerful tool to refine existing approaches and to design new experimental questions and methodologies.

A particular type of migraine with CSD is familial hemiplegic migraine type 3 (FHM3). It is caused by a mutation in the gene known as *SCN1A* (Castro et al. 2009; Shao et al. 2018). *SCN1A* codes for the voltage-gated sodium channel Na_V_1.1. This sodium channel controls the excitability of inhibitory neurons (Yu et al. 2006). The *SCN1A* mutation induces a gain of function in Na_V_1.1, increasing the excitability of inhibitory neurons (Bertelli et al. 2018; Cestèle et al. 2008, 2013). This hyperexcitability of inhibitory neurons is considered to be the origin of CSD in FHM3 (Cestèle et al. 2008, 2013).

The link between inhibitory-neuron hyperexcitability in FHM3 and CSD has been explored in several studies (Cestèle et al. 2008, 2013; Dhifallah et al. 2018). The prevailing hypothesis is that hyperactive inhibitory neurons rapidly increase extracellular K^+^. This increases overall neural excitability, including in excitatory cells and further boosting extracellular K^+^. This positive feedback perturbs ionic homeostasis, leading to a sustained neural depolarization. This hypothesis has been studied both experimentally (Auffenberg et al. 2021; Cestèle et al. 2008, 2013; Chever et al. 2021; Dhifallah et al. 2018) and computationally (Baspinar et al. 2025; Desroches et al. 2019; Lemaire et al. 2021), with a particular focus on the role of inhibitory neurons in CSD.

Previous computational CSD models fall into four categories: cellular, diffusion, network, and neural field models.

At cellular level, the first model linking gain of function in Na_V_1.1 and CSD initiation was proposed in (Desroches et al. 2019), based on a coupled pair of excitatory and inhibitory neurons. This coupling takes into account the ionic currents linked to extracellular K^+^. This model was further developed in (Chever et al. 2021) to reproduce experimental data obtained *in vitro* from mouse brain slices, and then in (Lemaire et al. 2021) with additional biophysical mechanisms and an extension to epileptiform activity. Noteworthy, these cellular models focus only on CSD initiation, not on its propagation.

Diffusion models focus on CSD propagation by considering the diffusion of ions and neurotransmitters involved in CSD. In (Tuckwell and Miura 1978), a system of reaction-diffusion equations were used to predict several qualitative properties of CSD propagation through an increase of extracellular K^+^ concentration. A similar framework was proposed in (Shapiro 2001), by including effects of cell volume and osmotically induced ion movements. One can note as well the model proposed in (Reggia and Montgomery 1996), exploring the link between CSD and hallucinations during migraine aura, in which CSD propagation was captured by a neural field type framework defined on a discrete tessellation of the cortex. Yet, diffusion models focus only on ions and neurotransmitters, and they do not represent any neural dynamics explicitly.

CSD network models represent neural dynamics explicitly through specific variables, each neuron of the network being modeled via a system of (often biophysical) differential equations. In (Florence et al. 2009), a three-neuron network was proposed to study the relation between the extracellular K^+^ concentration and CSD propagation. Larger networks were considered in (Conte et al. 2018; Huguet et al. 2016), by considering also the role of astrocytes in CSD dynamics. Finally, CSD was studied from a multiple-timescale perspective by using a large network setting in (Reyner-Parra et al. 2023).

Neural fields are formulated as coarse-grained continuum limits, providing a computationally efficient approximation of network dynamics, while preserving their essential collective behavior. Consequently, neural fields enable the study of CSD propagation in large-scale biological circuits and motivated our choice of a neural field with a pair of excitatory and inhibitory populations in (Baspinar et al. 2025).

An important limitation of most of the abovementioned models is that they only focus on neurons. They do not take into account astrocytes, which are considered to have important roles in CSD (Eren-Koçak and Dalkara 2021; Yang et al. 2024). Astrocytes are glial cells that regulate the ionic homeostasis in the extracellular matrix (Eren-Koçak and Dalkara 2021). Specifically, they clear excess K^+^ from the extracellular matrix. Therefore, they are closely linked to CSD propagation (ErenKoçak and Dalkara 2021; Yuan et al. 2019). This is the motivation for us to study the role of astrocytes in CSD propagation.

Our model is an extension from the neural population model proposed in (Baspinar et al. 2025). In the neural population model, excitatory and inhibitory population activity is represented in terms of average membrane potential of each population. The potential variables are coupled to a third state variable, which represents the extracellular K^+^ concentration. This model reproduces successfully the CSD propagation dynamics represented in terms of propagation speed. We extend this model by introducing a fourth state variable representing the activity-dependent recruitment of astrocytic K^+^ uptake, mediated by mechanisms such as Na^+^*/*K^+^-ATPase and Kir channel activity. This is a phenomenological variable representing the activation level of mechanisms involved in K^+^ uptake, considering uptake in isolation from other astrocytic functions.

An important novelty of the extended model is the inclusion of extracellular K^+^ clearance from the extracellular matrix. This clearance is modeled by a nonlinear term in the extracellular K^+^ equation, coupled to astrocyte population activity. The astrocytic dynamics follow the same formalism as the neural model in (Baspinar et al. 2025). Analogous to the neural population transfer function, the astrocyte transfer function has three regimes determined by the extracellular K^+^ concentration, allowing the model to capture how K^+^ levels modulate astrocytic K^+^ clearance. All these novelties provide a more complete framework and better fit of experimental data, compared to the neural population model in (Baspinar et al. 2025). See Section 2 and Appendix A for details.

Overall, the goal of these models is to reproduce the network-level dynamics, rather than the biophysical properties of individual neurons. In the present framework, excitatory and inhibitory neurons are represented as two populations, and the model focuses on their collective interactions and on the dynamics of extracellular K^+^ and astrocytic buffering.

One could ask about the relevance of modeling astrocytes as a population. Indeed, representing astrocytes as an additional population is possibly not the only way for such an extension towards astrocytic dynamics. Alternatively, one could introduce a function in the equation describing the extracellular K^+^ concentration, which would capture the K^+^ clearance from the extracellular matrix performed by astrocytes. However, this does not allow to represent explicitly astrocytic activity. Therefore, we cannot take into account the effect of neural firing rate on the astrocytic activation and, as a result, the interactions between neurons and astrocytes cannot be represented explicitly.

Another reason for considering the astrocytes as an additional population is that it allows us to link certain parameters to the experimental conditions which can be controlled optogenetically, thereby providing model-derived insights linked to experimental conditions.

The present work builds upon the neural population model proposed in (Baspinar et al. 2025) in three ways. First, it extends the neural dynamics to astrocytic dynamics. Second, it takes into account the astrocytic K^+^ clearance from the extracellular matrix, which is coupled to the astrocytic dynamics. Third, the model parameters are linked to pharmacological and/or optogenetic conditions modulating not only neural dynamics but also astrocytic dynamics. This provides model-derived insights regarding the role of astrocytes in FHM3-related CSD. Indeed, we show that including astrocytic dynamics provides a better fit to experimental data, and demonstrate that astrocytes have the potential to slow down and stop the CSD propagation as the inhibitory neural activity increases. The Matlab code used in this paper is available in (Baspinar et al. 2026).

## 2 Model

We introduce the astro-neural-field model that explicitly couples excitatory and inhibitory neural populations with astrocytic dynamics. This model accounts for astrocytic activity-dependent extracellular K^+^ regulation, thereby capturing the interactions between neurons and astrocytes that are critical for understanding CSD mechanisms in FHM3.

In the model, variables *v*_e_ (resp. *v*_i_) denote the average membrane potential of the excitatory (resp. inhibitory) neural population, *r*_a_ is a phenomenological variable representing activity-dependent recruitment of astrocytic K^+^ uptake such as Na^+^*/*K^+^ − ATPase and Kir channels, and *c*_*K*_ is the extracellular K^+^ concentration. The model is given by the following integro-differential system:

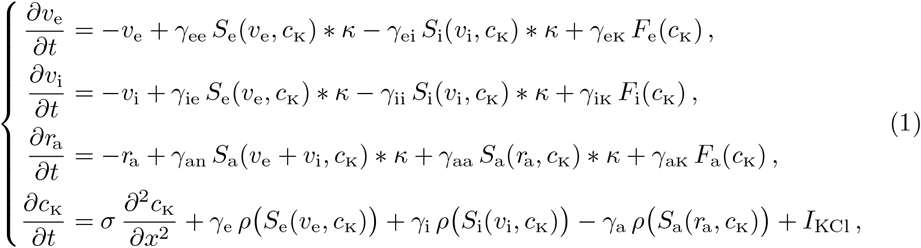

for all 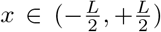, *t >* 0. All the weighting coefficients *γ*_◊ ♦_ and *γ*_◊_ are positive and serve to modulate the contribution of their associated components (for ◊ and ♦ corresponding to e, i, a, n or K, where n is the index for any neural population). The diffusion coefficient *σ* is strictly positive. See the below subsections for the model components and see Appendices A and B for further elements regarding the modeling rationale and discretization.

Throughout this article, we will consider system (1) with periodic boundary conditions, i.e. 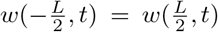 for all *t* ≥ 0 (*w* = *v*_e_, *v*_i_, *r*_a_, *c*_*K*_), and take initial conditions 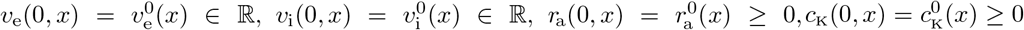. Note that, due to the initial conditions and the structure of system (1), it is straightforward to verify that the components *r*_a_(*x, t*) and *c*_K_(*x, t*) remain non-negative for all *x* and *t*.

The equation of astrocyte variable *r*_a_ follows the same population-level structure as the neural-field equations because it describes the evolution of a variable driven by external inputs and intrinsic interactions. The linear negative feedback term − *r*_a_ represents relaxation of astrocytic recruitment toward baseline in the absence of stimulation. The term *γ*_an_ *S*_a_(*v*_e_ + *v*_i_, *c*_K_) *∗ κ* models activation of astrocytes by surrounding neural activity through neuron-to-astrocyte signaling mechanisms. The term *γ*_aa_ *S*_a_(*r*_a_, *c*_K_) *∗ κ* represents the propagation and reinforcement of astrocytic activity through inter-astrocyte communication, such as gap-junction-mediated signaling. Finally, *γ*_aK_ *F*_a_(*c*_K_) captures the direct recruitment of K^+^-uptake mechanisms by elevated extracellular K^+^. Together, these terms provide a phenomenological population-level description of activity-dependent astrocytic K^+^ regulation.

Overall, system (1) couples the dynamics of *v*_e,i_ and *r*_a_ via non-local interaction terms (*S*_◊_ (,) *∗ κ*) modeling spatially distributed synaptic inputs to the neural populations, and the inputs to the astrocyte population, which are mediated by neurotransmitter signaling as well as by gap-junction coupling. Moreover, the dynamics of *v*_e,i_ and *r*_a_ are coupled with a diffusive K^+^ field through nonlinear transfer, activation (*F*_◊_ (*c*_K_)), and source (*γ*_◊_*ρ*(*S*_◊_)) functions. All these couplings allow us to generate synchronized CSD activity patterns; see Fig. 2. Finally, *I*_KCl_ is a nonlinear function that locally decays in space and time. It represents the KCl drops used in the experiments to provoke CSD in cortical tissue.

**Fig. 2.**
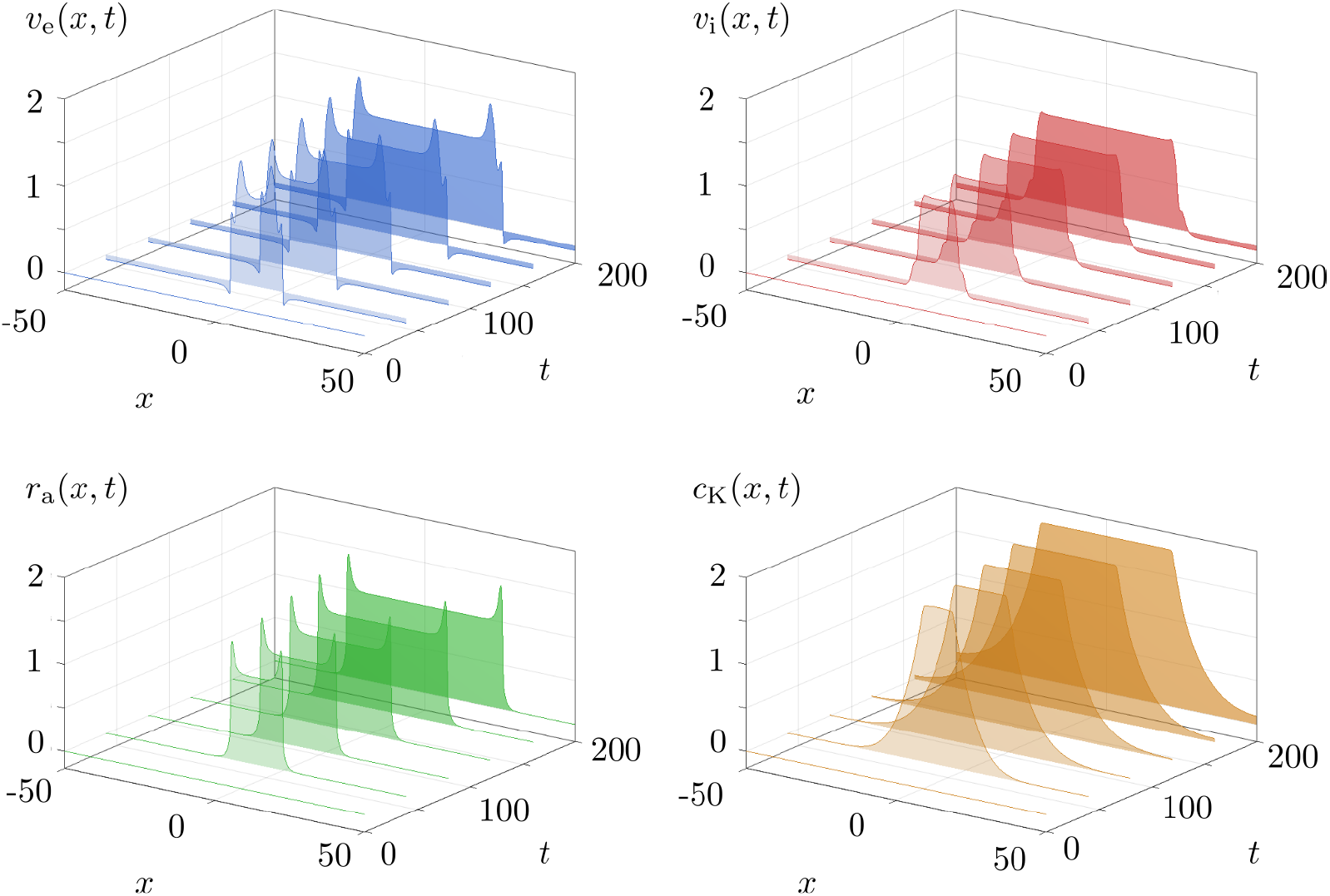
Simulation of system (1) with control parameters. Time integration of model variables is sampled at *t* = 0, 40, 80, 120, 160, 200. CSD is induced by the KCl stimulus input applied at *t* = 2. CSD activity patterns propagate symmetrically from the center (*x* = 0). Parameter values are given in Table A1.

**Fig. 3.**
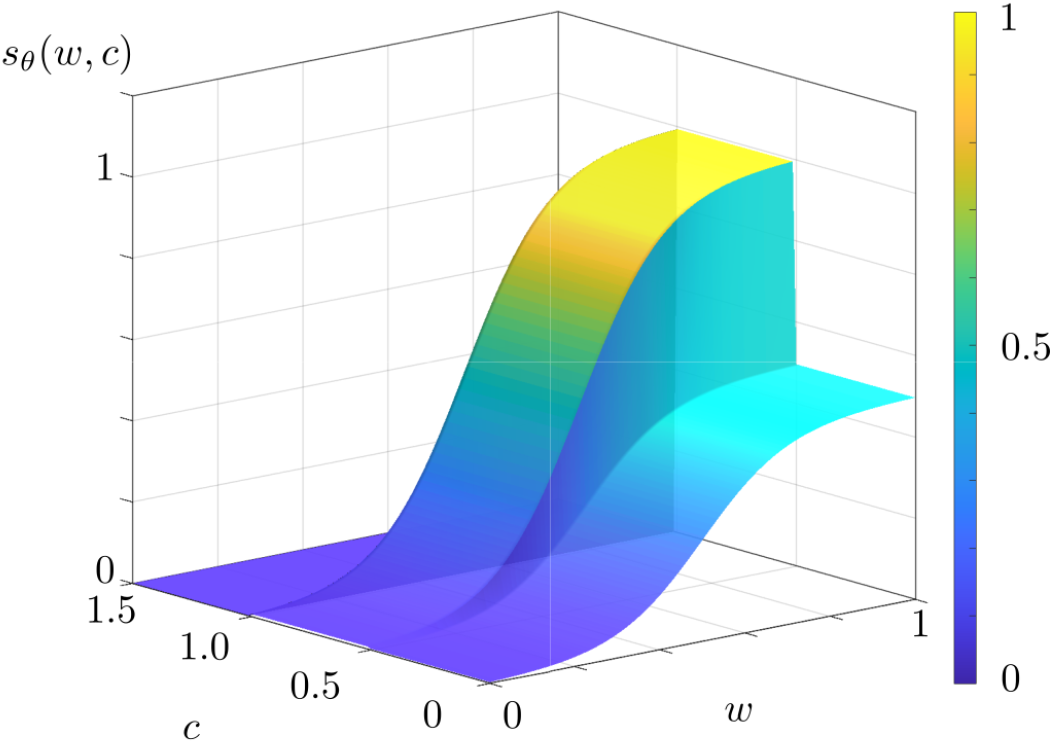
Population transfer function. Function *s*_*θ*_(*w, c*) is plotted with respect to *w* and *c*, where *θ*^1^ = 0.5, *θ*^2^ = 1.0, *θ*^3^ = 10, *θ*^4^ = 0.5.

If the astrocyte population is removed and the term describing astrocytic activity-dependent K^+^ clearance is omitted from the diffusive K^+^ field, system (1) reduces to the neural field model proposed in (Baspinar et al. 2025). This correspondence highlights that the present model can be viewed as a natural extension of the earlier neural field formulation, explicitly incorporating astrocyte–neuron interactions.

In the following, we explain mathematical structures of the model components. We refer to Appendix A for the rationale behind these components.

### Population transfer functions *S*_*◊*_ (*w, c*)

The so-called population transfer functions *S*_*◊*_ (*w, c*) are of the form:

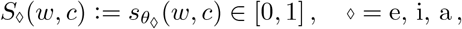

for some given function *s*_*θ*_(*w, c*) of *w*, which could be *v*_e_, *v*_i_, *r*_a_ or *v*_e_ + *v*_i_, and of a given K^+^ concentration *c*. Here *s*_*θ*_(*w, c*) is parametrized by a 4-dimensional parameter *θ* = (*θ*^1^, *θ*^2^, *θ*^3^, *θ*^4^) depending on the type of population, e, i or a:

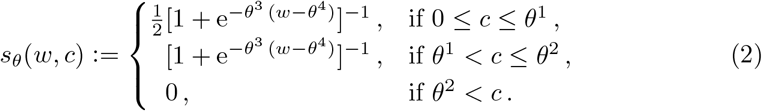

The population transfer functions *S*_*◊*_ (*w, c*) are interpreted as:

(*i*) for _◊_ = e, i, *S*_e_(*v*_e_, *c*) [resp. *S*_i_(*v*_i_, *c*)] is the proportion of spiking excitatory [resp. inhibitory] neural population for a given voltage *v*_e_ [resp. *v*_i_], at a given K^+^ concentration *c*;

(*ii*) for _◊_ = a:

- *S*_a_(*v*_e_ + *v*_i_, *c*) is the proportion of the astrocyte population that is activated for recruitment of astrocytic K^+^ uptake for a given voltage *v*_e_ + *v*_i_, at a given K^+^ concentration *c*;
- *S*_a_(*r*_a_, *c*) is the proportion of the astrocyte population that activated for recruitment of astrocytic K^+^ uptake according to the recruitment level *r*_a_, at a given K^+^ concentration *c*;

in all cases, *S*_◊_(*w, c*) represents a proportion of a population and therefore belongs to [0, 1].

In the first three equations of (1), the population transfer functions appear in a non-local integral form, expressed as convolution products:

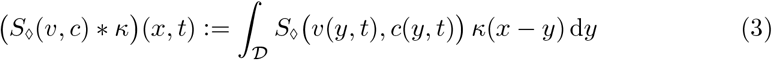

where *κ*(*x*) denotes the connectivity kernel, which describes how the strength of interactions between cells decays with spatial distance. In this work, we adopt a short-range excitatory kernel of the form:

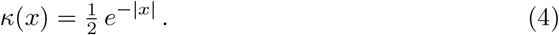

The spatial convolution of the population transfer functions with the connectivity kernel enables the emergence of spatially propagating activity patterns.

### K^+^-dependent activation functions *F*_◊_(*c*)

The term *F*_◊_(*c*) models the K^+^-dependent modulation of population activity. It is represented by a logistic activation function:

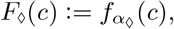

with

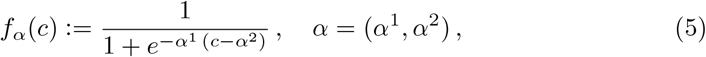

where *α*^1^ controls the sensitivity to changes in *c, α*^2^ determines the concentration threshold at which the response becomes half-maximal.

### K^+^ source function *ρ*(*s*)

The function *ρ*(*s*) describes the contribution of the K^+^ source generated by the neural or astrocytic populations. It is modeled as a bell-shaped, symmetric activation profile:

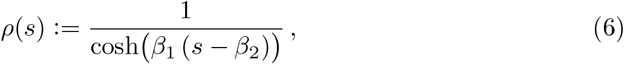

where *β*_1_ controls steepness, and *β*_2_ specifies the activation threshold. The function *ρ* is parameterized so that the input values *s* lie within the left half of its domain. Correspondingly, the outputs of *ρ* correspond to its increasing segment, i.e. the region of the curve that lies to the left of its peak.

### KCl stimulus input *I*_**KCl**_(*x, t*)

Finally, in the experimental setting, CSD is initiated by applying potassium chloride (KCl) drops onto the cortical surface. This induces a rapid and pronounced increase of the extracellular K^+^ concentration in a small cortical region, thereby triggering the onset of CSD at that location. In the model, this manipulation is represented as a localized external input *I*_KCl_ entering the K^+^-concentration equation in (1). We prescribe

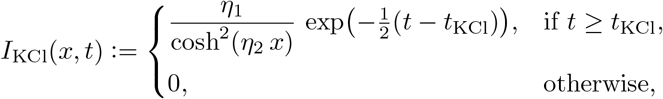

where *t*_KCl_ ∈ (0, *T*) denotes the time at which the KCl drops are applied. The input is localized both in space and time: the factor cosh^−2^(*η*_2_ *x*) produces a smooth and symmetric spatial bump centered at *x* = 0, taken as the application site, with spatial decay rate controlled by *η*_2_ *>* 0. The temporal term 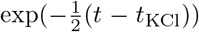 models the rapid dispersion of KCl after its application. The parameter *η*_1_ *>* 0 sets the peak amplitude of the stimulus.

## 3 CSD propagation in experiments

We verify our model via a comparison of our simulation results to the experimental data presented in (Baspinar et al. 2025). This data was obtained from mouse brain slices to study CSD in terms of its propagation speed, with a particular focus on the role of the inhibitory neurons.

Inhibitory neurons may exert two opposing effects in a network of neurons: one via classical inhibitory synaptic transmission, which hyperpolarizes and suppresses target cells, and the other via activity-dependent increases in extracellular K^+^, which can paradoxically depolarize and silence the network. The experiments in (Baspinar et al. 2025) study these effects by selectively enhancing or blocking synaptic inhibition, combined with optogenetic control of neural activity.

Increase or blocking of inhibitory synaptic transmission is achieved by perfusing in the brain slices the following pharmacological agents: isoguvacine (ISO) and gabazine (GBZ). ISO enhances inhibitory synaptic transmission. On the contrary, GBZ blocks it. These pharmacological interventions are modeled by varying *γ*_ei_ and *γ*_ii_ in the model.

Optogenetic modulation involves exposing the brain slices to blue light. In slices from genetically modified lines, this light activates the channelrhodopsin-2 ion channel (ChR2), increasing both inhibitory neural activity and inhibitory synaptic transmission. In the model, the elevated inhibitory activity is represented by increasing *γ*_i_ to reflect higher extracellular K^+^ release to the extracelluar matrix, while the enhanced inhibitory synaptic transmission is captured by increasing simultaneously *γ*_ei_ and *γ*_ii_.

CSD was triggered by focal KCl application to brain slices, and its propagation was measured with intrinsic optical signal (IOS) imaging and local field potential recordings. Each slice underwent a single CSD event, tracked through a sequence of images.

Propagation speed was calculated from wavefront displacement across four successive frames, producing three measurements that were averaged for a slice specific value. Slice-wise speeds were then averaged to yield the experimental propagation speed, which we denote by *v*_obs_. Local field potential recordings independently confirmed these values by computing speed from the time lag Δ*t* between wavefront passage at two electrodes. The electrode positions on the brain slice are denoted by *x*_1_ and *x*_2_ (Fig. 1), and the distance between them is *d* = |*x*_2_ − *x*_1_|, yielding the propagation speed *d/*Δ*t*.

Control case refers to the CSD propagation in the absence of any pharmacological or optogenetic intervention in the experiments. The control value of the propagation speed *v*_obs_ is 2.2 mm/min. In addition to the control case, four experimental conditions are considered (Fig. 4a):

**Fig. 4.**
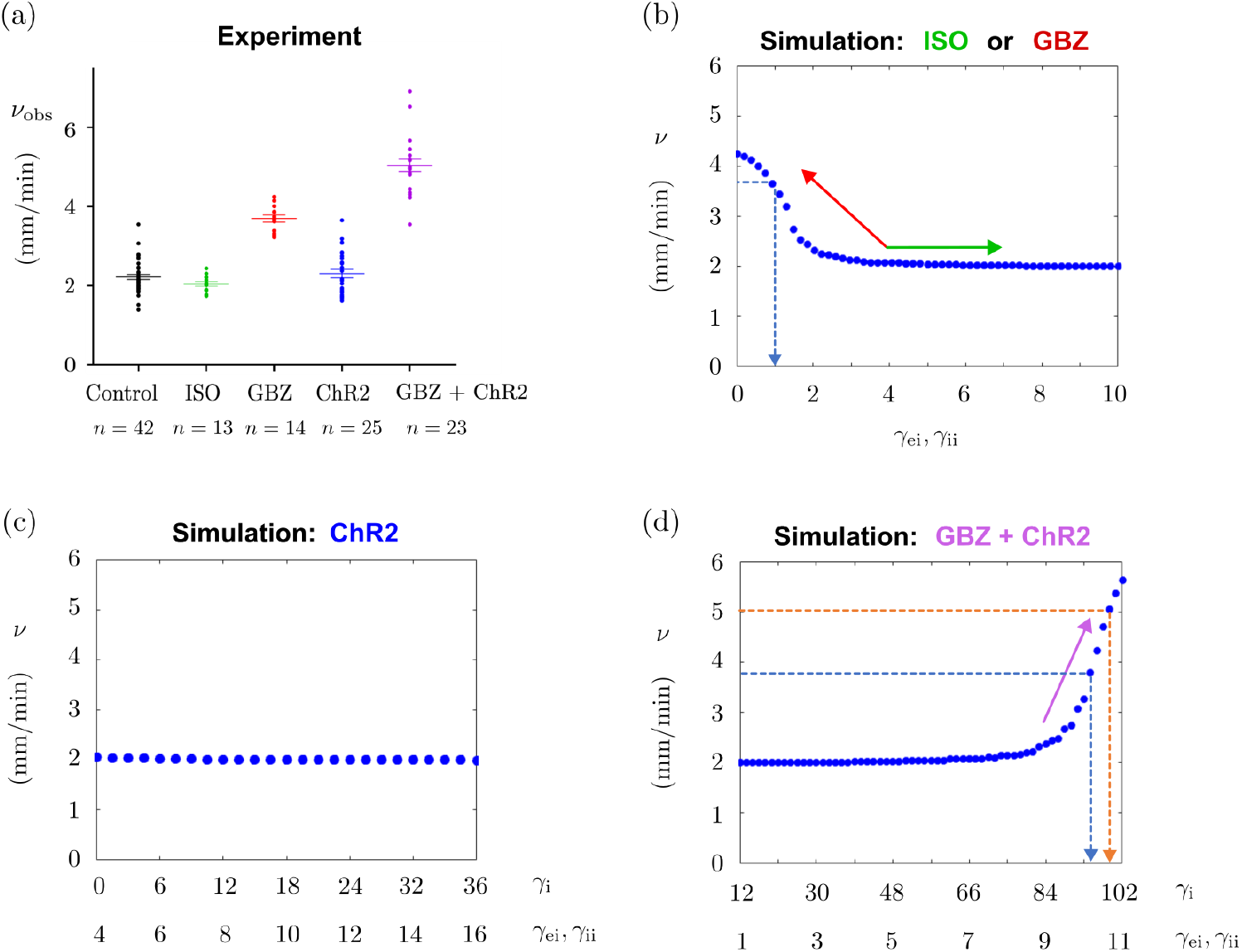
Experimental and simulation results of CSD propagation speed. (a) Experimental results from (Baspinar et al. 2025). CSD propagation speeds recorded in the control case and in (EC1)-(EC4). The propagation speed is denoted by *v*_obs_. Experimental conditions are denoted on the horizontal axis. Long horizontal bars show the mean value of the propagation speed and the short horizontal bars denote the standard deviation. They are obtained from measurements averaged over the number of slices denoted by *n* below the horizontal axis. Propagation speed obtained from each slice is represented with dots. In (b), (c) and (d), propagation speed *v* is computed from the model simulations by using (C2) from Appendix C. Each blue dot denotes the *v* value corresponding to the parameter given in the horizontal axis. (b) Simulation results corresponding to (EC1) and (EC2). ISO and GBZ effects are highlighted by green and red arrows, respectively. Dashed blue arrow indicates *γ*_ei_, *γ*_ii_ values matching the experimental speed *v*_obs_ in (EC2). (c) Simulation results corresponding to (EC3). (d) Simulation results corresponding to (EC4). GBZ+ChR2 effect is highlighted by the purple arrow. Dashed blue and orange arrows indicate the experimental speed *v*_obs_ values obtained in (EC1) (3.8 mm/min) and (EC4) (5 mm/min), respectively. In all plots, only the parameters indicated in the horizontal axes are varied. The rest of the parameters have the control values given in Table A1.

(EC1) ISO: Enhancement of inhibitory synaptic transmission. No significant change is observed in the propagation speed compared to the control value.

(EC2) GBZ: Blocking of inhibitory synaptic transmission. An increase to 3.8 mm/min is observed in *v*_obs_.

(EC3) ChR2: Enhancement of both inhibitory neural activity and inhibitory synaptic transmission. No significant change is observed in *v*_obs_.

(EC4) GBZ+ChR2: Enhancement of inhibitory neural activity and blocking of inhibitory synaptic transmission. Application of GBZ and activation of ChR2 via optogenetic illumination increases *v*_obs_ to 5 mm/min. This increase is higher compared to the increase in (EC2), where only GBZ is applied.

Corresponding simulation results of the model are given in Fig. 4b, c and d, which we will discuss in the following section.

## 4 Results

### 4.1 Better fit of experimental data

The astro-neural model quantitatively reproduces the experimentally observed CSD propagation speeds across all conditions (EC1)-(EC4). To assess this, we computed the propagation speed as described in Appendix C. Control parameters (Table A1) were selected to match the mean propagation speed measured experimentally under control conditions, namely 2.2 mm/min (Baspinar et al. 2025); see Fig. 4.

Fig. 4b shows the simulation results corresponding to cases (EC1) and (EC2). In (EC1), ISO enhances inhibitory synaptic transmission, which is modeled by increasing the connectivity weights *γ*_ei_, *γ*_ii_ from their control value. Experimentally, ISO does not produce a noticeable change in propagation speed, and this behavior is accurately reproduced by the model, as illustrated by the plateau highlighted by the green arrow in Fig. 4b.

In (EC2), GBZ blocks inhibitory synaptic transmission, leading experimentally to an increase in propagation speed to approximately 3.7 mm/min. In the model, this condition is captured by decreasing *γ*_ei_, *γ*_ii_ from their control value. The resulting increase in propagation speed closely matches the experimental observation, as indicated by the blue dashed arrow in Fig. 4b.

Importantly, the present astro-neural model resolves a key limitation of the previous neural field framework (Baspinar et al. 2025). In that model, complete suppression of GABAergic transmission (*γ*_ei_, *γ*_ii_ = 0) produced unrealistically large propagation speeds (> 10 mm/min; see (Baspinar et al. 2025, Fig. 6B)), far exceeding experimental values. In contrast, the astro-neural model yields a saturation around *v* = 4.2 mm/min under the same conditions, which is substantially closer to the experimentally observed value. This demonstrates an improved quantitative consistency with experimental data.

This improvement is due to the introduction of astrocytes in the model. As inhibitory synaptic interactions are turned off, excitation that the neural populations go through are not balanced. This results in an abrupt increase in the K^+^ release to the extracellular matrix via *γ*_e_ *ρ*(*S*_e_) and *γ*_i_ *ρ*(*S*_i_). In the absence of any K^+^ uptake, as in the case of the previous model (Baspinar et al. 2025), this results in lack of control in the diffusion speed of increased K^+^, inducing an increase in CSD propagation speed. By including the K^+^ uptake term −*γ*_a_ *ρ* (*S*_a_) driven by the dynamic variable *r*_a_, we can compensate the increase in extracellular K^+^ arising from increased neural activity due to the turning off the inhibitory synaptic interactions. This allows us to provide a better control on the propagation speed of CSD as the synaptic interactions vanish.

Fig. 4c shows the results for condition (EC3), where optogenetic activation of ChR2 enhances both inhibitory neural activity and synaptic transmission. This is modeled by increasing *γ*_i_ together with *γ*_ei_, *γ*_ii_. Experimentally, the propagation speed remains close to the control value, and this behavior is reproduced by the model, with simulated speeds remaining near 2.2 mm/min.

Fig. 4d presents the results for condition (EC4), where GBZ and optogenetic activation are applied simultaneously. Experimentally, this leads to a marked increase in propagation speed to approximately 5 mm/min. The model reproduces this increase when *γ*_ei_, *γ*_ii_ and *γ*_i_ are varied accordingly from their control values, as highlighted by the orange dashed arrow in Fig. 4d.

Overall, these results demonstrate that the astro-neural model reproduces the full set of experimental conditions (EC1)-(EC4), while improving quantitative agreement in regimes where the previous model in (Baspinar et al. 2025) exhibited certain discrepancies regarding (EC2).

Finally, we provide in Fig. 5 a measure of sensitivity of the model to the fitting parameters *γ*_ei_, *γ*_ii_ and *γ*_i_, as well as to the parameters *γ*_ie_, *γ*_ee_, *γ*_an_, *γ*_aa_, *γ*_e_, *γ*_a_. We perturb these parameters around their control values, with a multiplicative noise whose amplitude is generated from standard normal distribution. More precisely, we simulate 50 trials of the model with control parameters. In each trial, we perturb the chosen parameter p *∈ {γ*_ei_, *γ*_ie_, *γ*_ee_, *γ*_ii_, *γ*_an_, *γ*_aa_, *γ*_e_, *γ*_i_, *γ*_a_*}* with a value *z* generated from standard normal distribution:

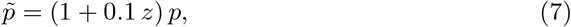

where 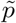 denotes the perturbed parameter. In this procedure, *z* is generated independently for each trial. Mean value and the standard deviation of the propagation speeds obtained from the trials with perturbed parameters are plotted in Fig. 5, separately for each perturbed parameter.

**Fig. 5.**
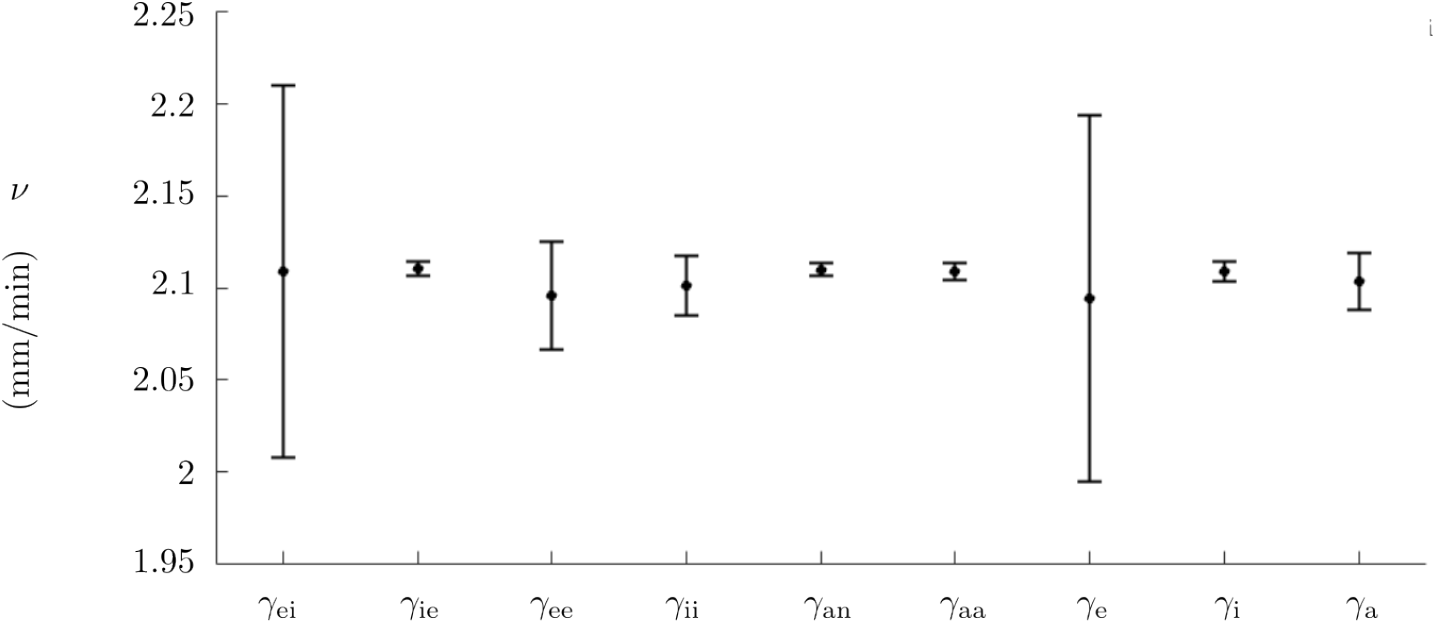
Sensivity analysis with respect to *γ*_ei_, *γ*_ie_, *γ*_ee_, *γ*_ii_, *γ*_an_, *γ*_aa_, *γ*_e_, *γ*_i_, *γ*_a_. Black dots denote the mean values obtained from 50 trials for each parameter given in the horizontal axes. Vertical lines show the standard deviations. See the text for more details.

### 4.2 Better control on CSD propagation

We time-stepped system (1) by using the control parameter set (Table A1). Fig. 2 shows a representative simulation in which all state variables are initialized at zero. An external input *I*_KCl_ is applied at *t* = 2, centered at *x* = 0, representing the experimental application of KCl used to trigger CSD.

The model generates waves that originate at the stimulation site and propagate symmetrically toward the boundaries at 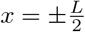. The propagating wavefront is characterized by elevated neural activity, followed by a sustained depolarization block. This block corresponds to moderately elevated membrane potentials in the neural populations. All state variables evolve in a synchronized manner during propagation; see Fig. 2.

This synchronization can be modulated by varying the astrocytic connectivity parameter *γ*_aa_, as shown in Fig. 6. The spatial convolution term *S*_a_(*r*_a_, *c*_K_) *∗ κ* effectively acts as a localized diffusion of astrocytic activity near the wavefront. Since astrocytes interact with neural populations primarily through extracellular K^+^, changes in astrocytic dynamics can alter propagation features without substantially modifying the underlying neural activity, provided *γ*_a_ remains fixed.

**Fig. 6.**
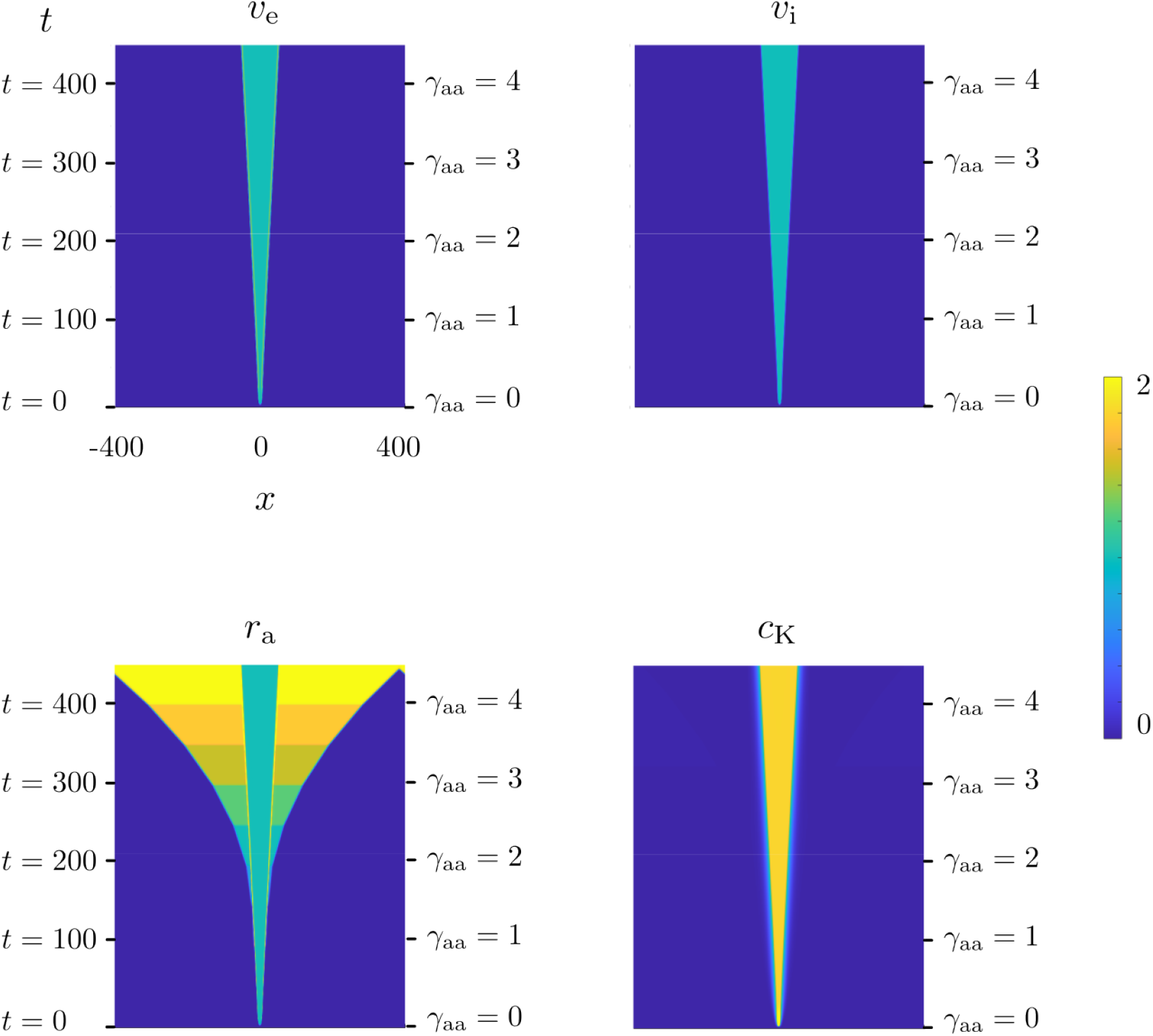
Space-time plots of the model variables where *γ*_aa_ is increased. Horizontal and vertical axes denote in all plots the space and time, respectively. The color map is the same for all plots and it denotes the value of corresponding model variable. In the simulations, *γ*_aa_ is increased from 0 to 4 by increments of 0.5 at the end of each time duration of 50, as highlighted in each plot. The rest of the parameters are the same as the control parameters given in Table A1. Here *L* = 800, *N* = 2^14^ and *T* = 450.

In addition, increasing *γ*_a_, which enhances K^+^ clearance from the extracellular space, leads to a progressive slowing of wave propagation and can ultimately suppress it; see Fig. 7. This illustrates how astrocytic regulation of extracellular K^+^ can modulate CSD dynamics within the model.

**Fig. 7.**
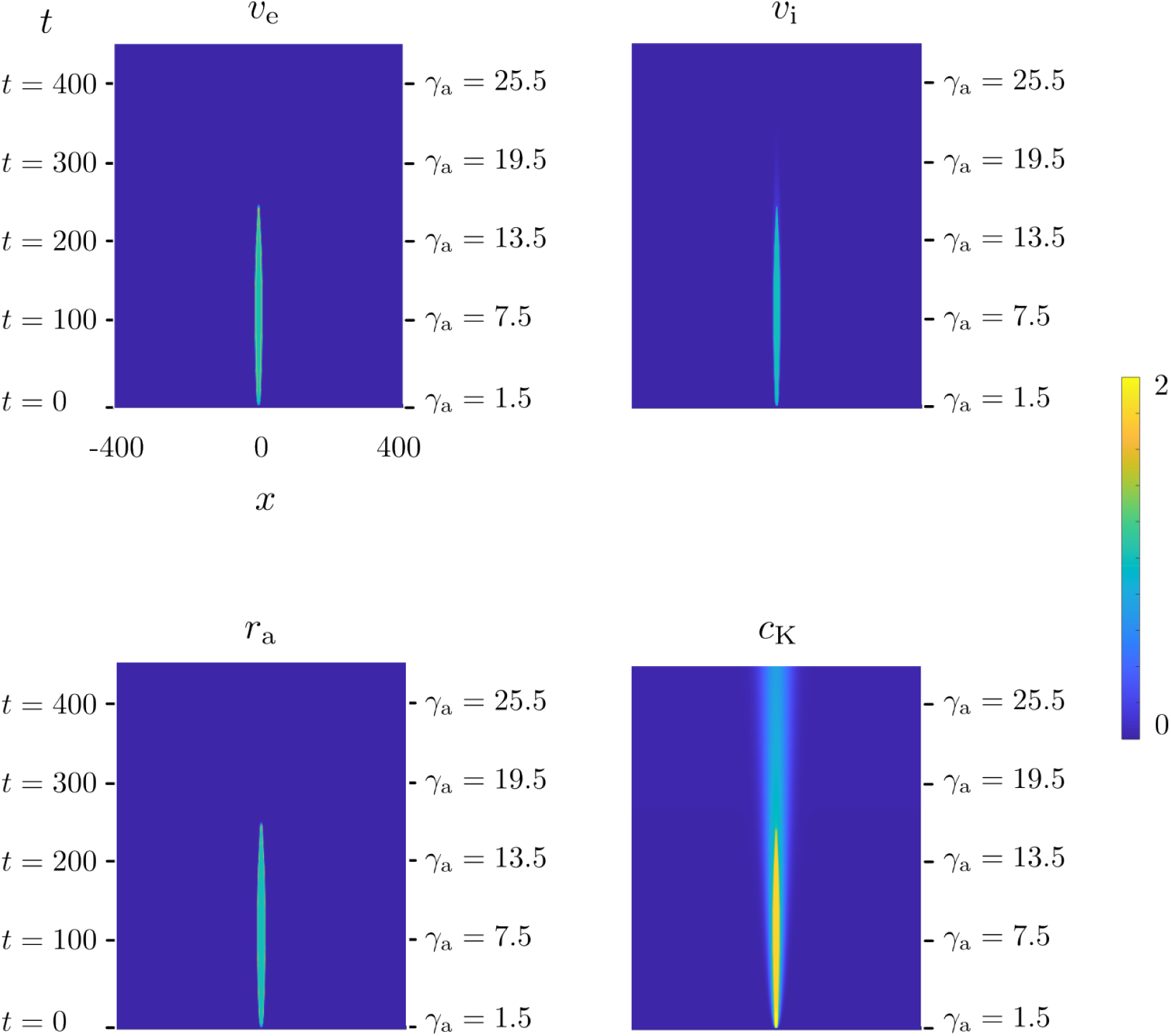
Space-time plots of the model variables where *γ*_a_ is increased. Horizontal and vertical axes denote in all plots the space and time, respectively. The color map is the same for all plots and it denotes the value of corresponding model variable. In the simulations, *γ*_a_ is increased from 1.5 to 25.5 by increments of 3 at the end of each time duration of 50. The rest of the parameters are the same as the control parameters given in Table A1. Here *L* = 800, *N* = 2^14^ and *T* = 450.

### 4.3 Model-derived insights

#### 4.3.1 Elevated astrocytic activity beyond CSD propagation

Astrocytic calcium waves have been reported to extend beyond the spatial range of CSD waves in brain slices (Charles and Brennan 2009; Peters et al. 2003). Within the model framework, this behavior can be explored by increasing the astrocytic connectivity parameter *γ*_aa_, which represents gap-junction coupling strength between astrocytes.

As shown in Fig. 6, increasing *γ*_aa_ enhances the spatial spread of astrocytic activity independently of the neural CSD wave. This allows the model to generate astrocytic activity that persists and propagates beyond the region affected by CSD (Fig. 8), consistent with experimental observations.

**Fig. 8.**
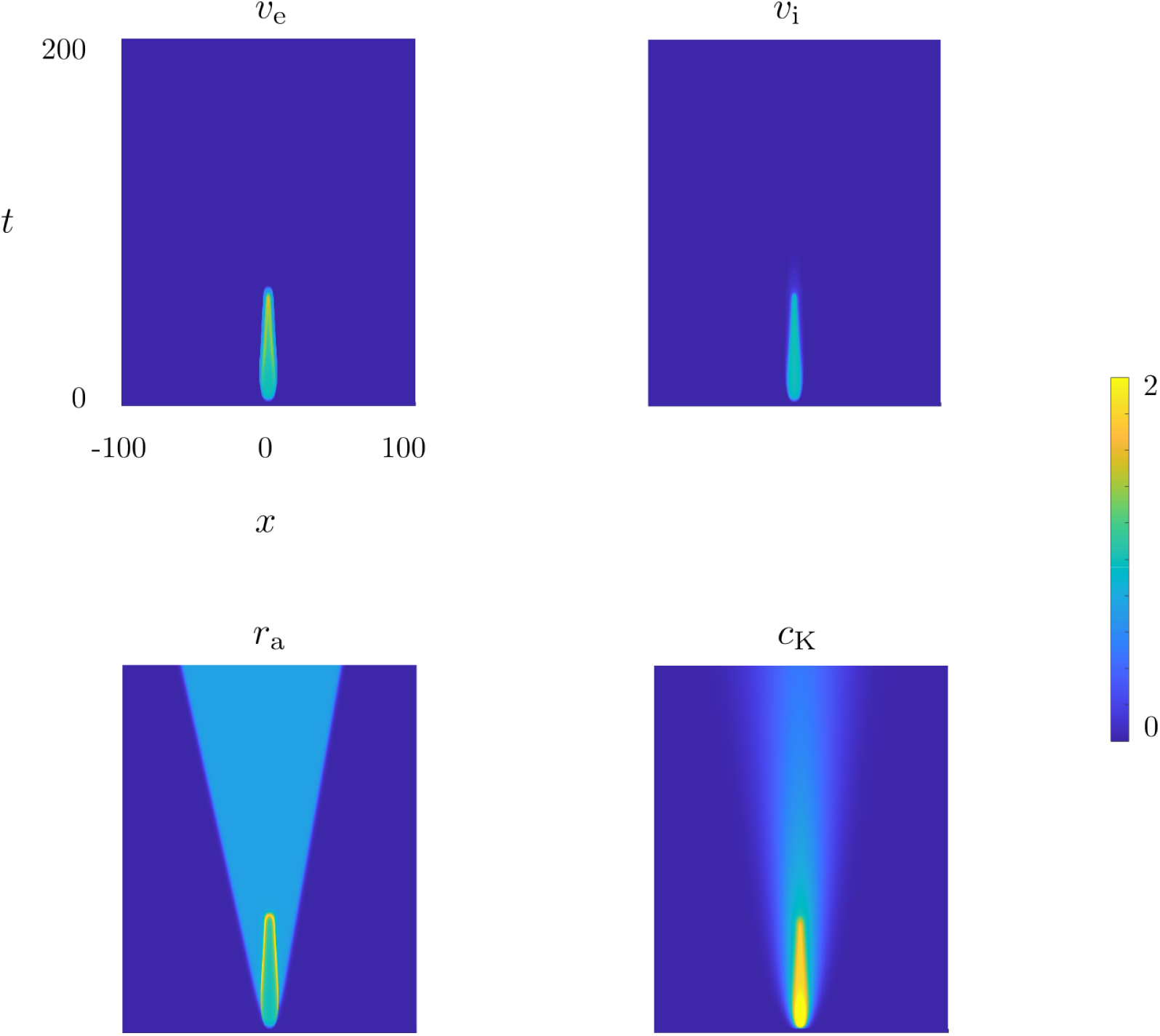
Space-time plots of the model variables where astrocytic activity propagation extends beyond CSD propagation. Horizontal and vertical axes denote in all plots the space and time, respectively. The color map is the same for all plots and it denotes the value of corresponding model variable. In the simulations, *γ*_aa_ = 1.5 and *γ*_a_ = 10.5. The rest of the parameters are the same as the control parameters given in Table A1. Here *L* = 200, *N* = 2^12^ and *T* = 200.

This behavior arises only when *γ*_aa_ exceeds a certain range; for lower values, astrocytic activity remains synchronized with neural dynamics (Fig. 2 and 7). This suggests that astrocytic waves are maintained by at least two distinct mechanisms: one dependent on CSD and another governed by gap-junction coupling between the astrocytes.

#### 4.3.2 Effect of astrocytic modulation on CSD propagation

Existing experimental and computational studies primarily focus on neural mechanisms (Baspinar et al. 2025; Desroches et al. 2019; Lemaire et al. 2021), while astrocytic contributions are typically not included, despite their established role in regulating extracellular K^+^ (Yang et al. 2024).

In the present model, astrocytic activity modulates extracellular K^+^ through the parameter *γ*_a_, which can be interpreted as representing optogenetic activation. Simulations show that increasing *γ*_a_ slows down CSD propagation and can eventually suppress it entirely; see Fig. 9.

**Fig. 9.**
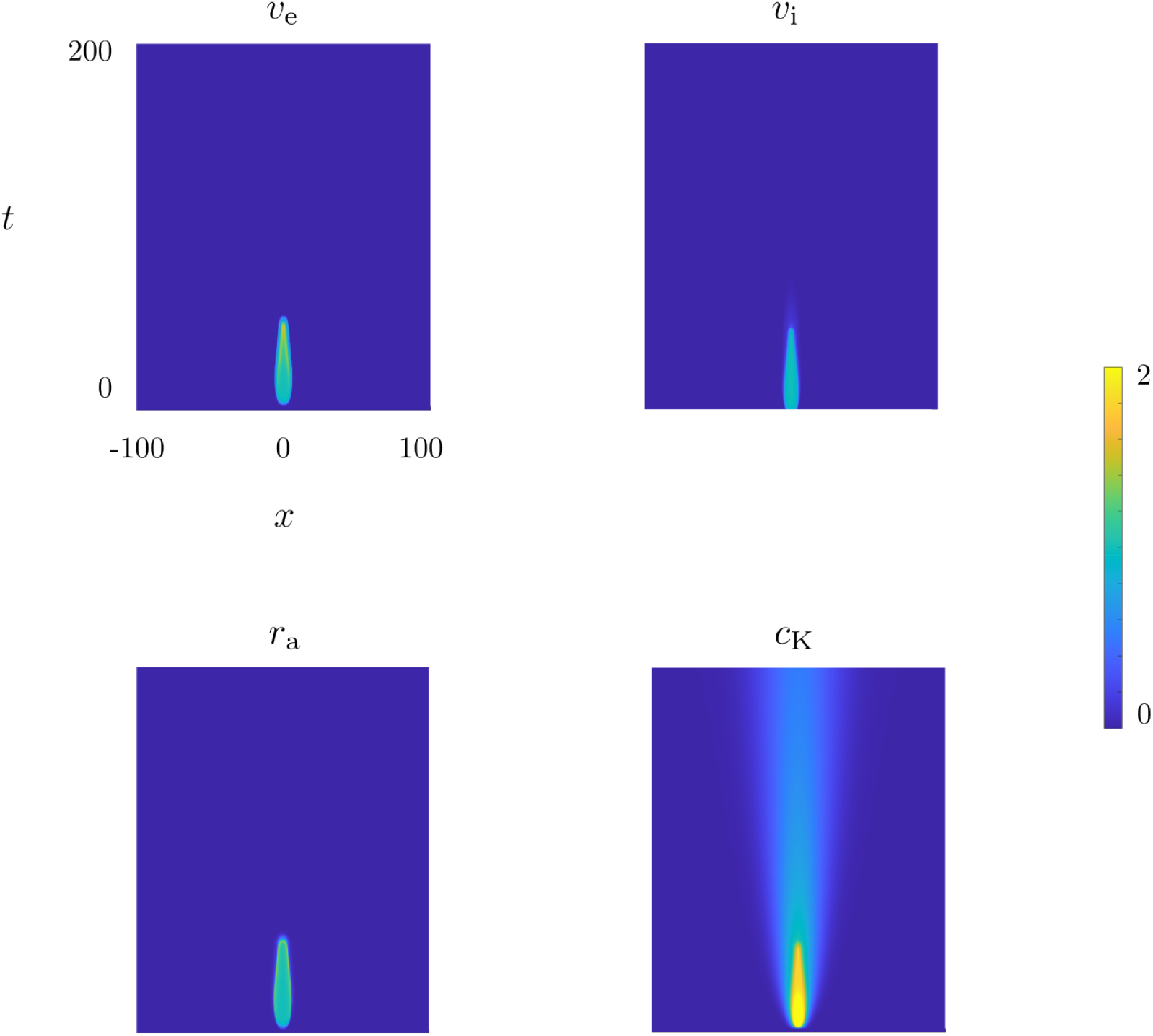
Space-time plots of the model variables where CSD propagation stops due to increased astrocytic activity of extracellular K^+^ clearance. Horizontal and vertical axes denote in all plots the space and time, respectively. The color map is the same for all plots and it denotes the value of corresponding model variable. In the simulations, *γ*_a_ = 13. The rest of the parameters are the same as the control parameters given in Table A1. Here *L* = 200, *N* = 2^12^ and *T* = 200.

These results indicate that astrocytic regulation of extracellular K^+^ can significantly influence CSD dynamics within the model. This effect is consistent with a modulatory role of astrocytes, complementing the primary contribution of inhibitory neural dysfunction in FHM3, studied in several works (Auffenberg et al. 2021; Cestèle et al. 2008; Chever et al. 2021; Desroches et al. 2019; Dhifallah et al. 2018; Lemaire et al. 2021).

## 5 Discussion

We presented a novel population model of neurons and astrocytes to study migraine-related CSD propagation. The model extends the neural field framework of (Baspinar et al. 2025) by incorporating astrocytes. In the previous model, excitatory and inhibitory neural populations are described by two state variables representing their average membrane potentials, coupled to a third state variable describing the extracellular K^+^ concentration. This coupling enables the investigation of how neural activity drives CSD through extracellular K^+^ accumulation, with particular emphasis on the facilitatory role of increased inhibitory activity. However, the previous model accounts only for contributions of neurons and neglects the potential influence of astrocytes on CSD propagation.

Astrocytes play a critical role in maintaining extracellular K^+^ homeostasis and are therefore thought to have significant roles in CSD propagation (Eren-Koçak and Dalkara 2021; Yuan et al. 2019). To account for this, we extended the previous neural field model by introducing a fourth state variable representing the activity-dependent recruitment of astrocytic K^+^ uptake. This variable is coupled to both neural population activity and extracellular K^+^ concentration, enabling us to investigate how pathological conditions affect astrocyte-mediated K^+^ clearance from the extracellular space in response to changes in neural activity.

The first reason for introducing astrocytic activity as an additional state variable is to enable its explicit tracking, similarly to (Moyse and Berry 2022). This allows us to examine the synchrony between increased neural activity and the activity-dependent recruitment of astrocytic K^+^ uptake during CSD propagation. In this way, the model provides an explicit representation of neuron-astrocyte interactions in the context of CSD dynamics. The second reason is that this formulation introduces an astrocyte-related parameter (*γ*_a_), which is directly associated with the optogenetic activation of ChR2 expressed in astrocytes, thereby enhancing astrocytic activity. This corresponds to one of the experimental conditions that can be controlled optogenetically. The third reason is that the inclusion of astrocytes allows us to incorporate into the extracellular K^+^ concentration equation a K^+^ uptake term that is dynamically driven by the astrocytic recruitment variable *r*_a_. This term compensates for the large increase in CSD propagation speed that occurs when inhibitory synaptic interactions are suppressed, thereby providing better control of the propagation speed than the previous model proposed in (Baspinar et al. 2025).

The model provides a better fit to the experimental data and enables better control over CSD propagation than the previous neural field framework (Baspinar et al. 2025). This, in turn, provides the following model-derived insights into FHM3-related CSD: (*i*) Elevated astrocytic activity can persist beyond the propagation phase, remaining active even after CSD has ceased. Within our model, this occurs when astrocyte-astrocyte interactions mediated by gap junctions are enhanced. (*ii*) Optogenetically enhanced astrocytic activity can suppress CSD propagation, similarly to the antiseizure effects of optogenetic stimulation of astrocytes reported in epilepsy (Zhao et al. 2022). According to our model, this effect can be achieved by increasing astrocytic activity through optogenetic activation of ChR2.

It is important to note that although astrocytes play an important role in maintaining extracellular K^+^ homeostasis and modulating CSD propagation, as discussed in (*ii*), they should not be regarded as a primary cause of FHM3 or of CSD initiation in this disorder. CSD susceptibility is influenced by many additional factors. Based on the current understanding of FHM3 pathophysiology, alterations in astrocytic activity are more appropriately interpreted as secondary responses that modulate disease progression rather than as primary pathological mechanisms, similarly to what has been reported in certain forms of epilepsy (Genin et al. 2026).

In our model, the population transfer functions are adaptive to the extracellular K^+^ concentration, as in the previous neural field framework (Baspinar et al. 2025). This adaptation is implemented through fixed threshold levels of extracellular K^+^, which define distinct regimes of cellular activity. Since these thresholds are time-invariant, the adaptation itself is not dynamic.

We adopt this simplified scheme instead of dynamic adaptation mechanisms (Kilpatrick and Bressloff 2010; Shaw and Kilpatrick 2024) because CSD dynamics are driven primarily by large elevations in extracellular K^+^, rather than by small and rapid fluctuations. These substantial increases establish distinct K^+^ regimes, within which cellular dynamics become relatively insensitive to minor variations in extracellular K^+^ concentration.

In the present work, our focus is on the propagation phase of CSD rather than the recovery phase after CSD. To better reflect the biological interpretation of astrocytic dynamics during CSD, we emphasize that the decrease of the astrocytic transfer function *S*_a_ at very high extracellular K^+^ concentrations should not be interpreted as a complete cessation of astrocyte activity or cell failure. Rather, it represents a phenomenological approximation of a transient saturation of astrocytic K^+^ buffering and clearance mechanisms under the extreme ionic load at the propagating wavefront of CSD. Experimental studies indicate that astrocytes remain metabolically active during CSD and can mobilize glycogen reserves to support energy-demanding processes (Feuerstein et al. 2016). Nevertheless, CSD is associated with a marked increase in energy consumption and a transient mismatch between metabolic demand and supply, accompanied by ATP depletion and impaired ionic homeostasis (Mies and Paschen 1984; Selman et al. 2004). Within this context, the decline of *S*_a_ is intended to capture a temporary saturation or insufficiency of astrocytic K^+^ regulation during propagation, rather than irreversible astrocytic dysfunction.

Astrocytes are classically viewed as buffers that clear excess extracellular K^+^. Normally, the mechanism of spatial buffering redistributes K^+^ within the astrocyte syncytium to maintain the K^+^ homeostasis, carrying the excess extracellular K^+^ towards extracellular regions with lower-K^+^ concentration. However, this redistribution has the potential to considerably raise extracellular K^+^ in the regions with lower-concentrations, effectively supporting the depolarization wave of CSD rather than suppressing it (Smith et al. 2006). This can be computationally studied in our model by modifying the K^+^ clearance term as a contributive term to the extracellular K^+^, coupled to the astrocytic activity variable.

FHM3-related CSD is thought to result from genetic mutations that increase the excitability of inhibitory neurons. Mutations in the same gene are also implicated in Dravet syndrome, a severe and drug-resistant form of epilepsy. In contrast to FHM3, however, these mutations are associated with reduced excitability of inhibitory neurons in Dravet syndrome (Aiba et al. 2023).

CSD is observed in both conditions and exhibits similar properties despite differences in the underlying mechanisms of initiation. Although our model is formulated for FHM3-related CSD, it can be readily adapted to epileptic CSD by increasing the activity threshold of the inhibitory population transfer function, thereby capturing the reduced inhibitory excitability characteristic of Dravet syndrome (Lemaire et al. 2025).

The use of a 1D spatial interval as the spatial domain of the model is both a limitation and an advantage. Limitation, because CSD is intrinsically a 2D cortical phenomenon and can exhibit complex spatiotemporal patterns, including retracting waves, spiral waves, localized wave segments, and stationary structures, depending on the underlying excitability and spatial coupling mechanisms (Dahlem and Isele 2013; Verisokin et al. 2017, 2023). Such dynamics cannot be captured in a 1D framework. Advantage, because the aim of the present work is not to investigate pattern formation in CSD, but rather to isolate and analyze the impact of astrocyte-mediated K^+^ regulation on the propagation of CSD. For this purpose, a 1D framework provides a useful and sufficient minimal setting in which the propagation mechanism can be studied without additional geometric effects arising from wave curvature, wave collisions, or spiral dynamics.

Importantly, the local mechanisms underlying our model, namely neuronal K^+^ release, extracellular K^+^ diffusion, and astrocytic K^+^ uptake, are independent of spatial dimensionality. Therefore, while a 2D extension would be expected to generate a richer set of spatiotemporal patterns, consistent with the previous studies (Dahlem and Isele 2013; Verisokin et al. 2017, 2023), we expect the qualitative role of astrocytic K^+^ regulation identified here to remain relevant. A detailed investigation of how these mechanisms interact with spiral waves, localized wave segments, or other complex 2D CSD patterns is an interesting direction for future work.

Finally, the model accurately reproduces propagation speeds in the range of 2− 5 mm/min, matching the experimental data from mouse brain slices (Baspinar et al. 2025). The model spans the speed range of CSD propagation in human, which is reported as approximately 3.5 mm/min in functional MRI studies of migraine aura (Hadjikhani et al. 2001). This supports the potential of the model to be developed towards a translational tool for understanding human migraine pathophysiology.

## Acknowledgements

This work has been supported in part by funding from the French government, through the France 2030 investment plan managed by the Agence Nationale de la Recherche (ANR), as part of the Université Côte d’Azur’s Initiative of Excellence (IdEx) Jedi (ANR-15-IDEX-01) and by the Laboratory of Excellence “Ion Channel Science and Therapeutics” (LabEx ICST ANR-11-LABX-0015-01, France) to MM. MM’s laboratory is a member of the “Fédération Hospitalo-Universitaire” InovPain (FHU-InovPain, France) and of the Université Côte d’Azur’s Strategic IdEx Program (PSI) “Ion Channels”.

## Appendix A

### Modeling rationale

The model describes the dynamics of excitatory and inhibitory neural populations through the mean activity variables *v*_e_ and *v*_i_, respectively. Interactions of these neural populations are crucial to cortical stability. The model uses the variable *r*_a_ to describe the astrocytic recruitment corresponding to K^+^ uptake. The astrocytic interactions with neurons and with other astrocytes are crucial to the K^+^ homeostasis in the extracellular matrix. Extracellular K^+^ accumulation is the principal driver of CSD (Pietrobon and Moskowitz 2014). Our model formalizes this by including the extracellular K^+^ concentration as a spatiotemporal variable *c* _*K*_. We provide below the detailed reasoning behind the choices of terms that describe the model variables *v*_e_, *v*_i_, *r*_a_ and *c*_K_.

### Population transfer functions *S*_◊_(*w, c*)

Population transfer function *S*_◊_ represents the mean output activity of the corresponding population _◊_ with respect to the cellular input that the population receives. In classical neural field models, it is typically chosen as a sigmoid function or one of its variants (Amari 1977; Wilson and Cowan 1972, 1973), as the sigmoid accurately captures the physiological characteristics of the neural input-output relationship (Albrecht and Hamilton 1982).

In contrast to the classical neural fields, the transfer functions in our model depends not only on cellular inputs *v*_e_, *v*_i_, *r*_a_ but also on extracellular K^+^ concentration *c* _*K*_, modeling the modulatory effects of the extracellular K^+^ on the neural excitability (Fig. 3). This is crucial to produce propagating waves with CSD characteristics.

The neural transfer function exhibits three K^+^-dependent regimes. Normal activity occurs at physiological extracellular K^+^ concentration. As K^+^ increases, a hyper-activity regime emerges, with high membrane potentials and synchronized spiking. This marks the onset and propagation of CSD. At extreme extracellular K^+^ levels, a depolarization block follows. The depolarization block suppresses neural spiking via moderate depolarization as it propagates.

These three regimes map quantitatively to the K^+^ levels defined in (2). The transfer function is a standard sigmoid (max=1/2) in the physiological regime (0 ≤ *c*_K_ ≤ *θ*^1^). In the hyperactivity regime (*θ*^1^ *< c*_K_ ≤ *θ*^2^), it becomes a high-gain sigmoid (max=1). Finally, in the depolarization block (*θ*^2^ *< c*_K_), the transfer function saturates at zero, representing suppressed neural population output (Fig. 3).

Astrocytic K^+^ clearance might go through a transient failure via three (possibly) interconnected mechanisms: (*i*) energy depletion caused by excessive ion pumping required to balance the extracellular K^+^ increase (van Putten et al. 2021), (*ii*) oxidative stress and inflammation from elevated astrocytic activity (Mogensen et al. 2021; Yang et al. 2024), and (*iii*) astrocyte endfoot swelling due to osmotic imbalance arising from intense absorption of extracellular K^+^, which physically reduces extracellular space and hinders clearance (Yang et al. 2024).

Our model implements this transient failure progression via dependence of astrocyte transfer function on the extracellular K^+^. For 0 ≤ *c*_*K*_ ≤ *θ*^1^ (physiological), a standard sigmoid (max=1/2) represents normal astrocytic function. For *θ*^1^ *< c*_K_ ≤ *θ*^2^ (high load), a high-gain sigmoid (max=1) models increased clearance effort. When *θ*^2^ *< c*_K_ (failure), the astrocyte population output vanishes, representing dysfunction from energy depletion, oxidative stress, and swelling. Astrocyte transfer function is, therefore, the same function as the one used for the neural populations. The only difference is that we use different parameters.

It is important to note that we consider the astrocytic recruitment corresponding to K^+^ uptake, i.e. ion clearance kinetics, in isolation. We do not take into account astrocytic activity related to enzymatic kinetics, or synaptic modulation. This has an important impact on our modeling choices: (*i*) K^+^ release to the extracellular matrix shares the same time scale as neural spiking, so does the K^+^ uptake. This is the reason that we use the same time scale for both neural and astrocytic activity. (*ii*) enzymatic kinetics are neglected, which otherwise would require using an asymmetric saturation profile as astrocyte transfer function. This is the reason that we use sigmoid functions as astrocytic population transfer functions in normal (*c*_*K*_ *< c*_*1*_) and high K^+^ (*c*_1_ *< c*_K_ *< c*_2_) regimes.

Choosing an asymmetric function (e.g. Hill-type) as the astrocyte population transfer function for the normal and high K^+^ regimes, instead of a sigmoid function, does not change noticeably the propagation speed. To verify it, we compared the propagation speeds obtained with the control parameters by using the sigmoid function defined in (2) and a Hill-type function given by

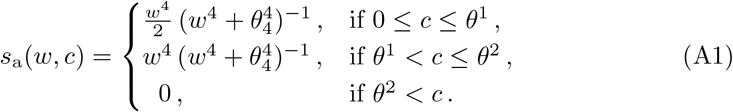

which gave 2.11 mm/min and 2.07 mm/min as propagation speeds, respectively.

The dependence of the astrocyte transfer function on extracellular K^+^ concentration is motivated by experimental evidence showing that glial K^+^-buffering mechanisms are strongly regulated by extracellular K^+^. In particular, the conductance of astrocytic Kir channels increases with extracellular K^+^ concentration, leading to enhanced K^+^ clearance when extracellular K^+^ rises (Kofuji and Newman 2004).

### Connectivity kernel *κ*(*x*)

Connectivity kernel *κ* models finite-range horizontal interactions between neurons, where these connections weaken with distance (Song et al. 2005). Astrocytes, which integrate local synaptic information and transmit it to neighboring astrocytes, are thought to exhibit activity decay over long distances, particularly if there is no regenerative coupling (Goldberg et al. 2010; Nowacka et al. 2025). Therefore, we choose *κ* as an exponentially decaying function of distance for both neural and astrocyte interactions.

Cellular interactions are modeled as the synaptic convolution given by (3). The spatial convolution is multiplied by the coefficient *γ*_◊_. This coefficient represents the strength of the connectivity from population to population _◊_. It is required because the strength of the horizontal connectivity or astrocytic influence need not necessarily be the same for every interacting population pair.

### K^+^-dependent activation functions *F*_◊_(*c*)

The dependence of astro-neural activity on the extracellular K^+^ concentration is reflected via a nonlinear mechanism linking the population dynamics to K^+^ processes. This interaction is represented by the K^+^-dependent activation function *F*_◊_ defined in (5).

In neural populations, elevated extracellular K^+^ pushes the neurons to an hyperactivity state, independently of the synaptic input. The activation functions *F*_◊_ abstract this effect into a phenomenological gain modulation of average membrane potential, capturing non-synaptic effects of K^+^ on the neural populations. Similarly, the function *F*_◊_ drives astrocytes toward a higher activity state as extracellular K^+^ rises. This reflects the increased K^+^ clearance demand.

We model the activation functions *F*_◊_ as sigmoids because extracellular K^+^ effects on neurons and astrocytes saturate beyond a certain K^+^ concentration in the extracellular matrix. These sigmoids are scaled by positive coefficients *γ*_eK_, *γ*_iK_, *γ*_aK_ to bridge the different quantities representing neural (membrane potential) and astrocytic (recruitment of astrocytic K^+^ uptake) activity.

### K^+^ source function *ρ*(*s*)

The source terms *ρ*(*S*_e_) and *ρ*(*S*_i_) represent activity-dependent release of K^+^ into the extracellular matrix from the excitatory and inhibitory neural populations, respectively. These terms phenomenologically abstract the K^+^ efflux to the extracellular matrix during action potentials and sustained depolarization. The term − *ρ*(*S*_a_) represents the activity-dependent clearance of K^+^ from the extracellular matrix by the astrocyte population.

In *ρ*(*S*_e_) and *ρ*(*S*_i_), the dependence of the function *ρ* on the outputs of the neural transfer functions ensures that (*i*) low neural firing produces weak K^+^ accumulation in the extracellular matrix, (*ii*) once the neural activity exceeds a critical level, K^+^ release to the extracellular matrix increases rapidly.

In − *ρ*(*S*_a_), the dependence on the output of the astrocyte transfer function provides the opposite: (*i*) low or moderate astrocytic activity clears normal quantities of K^+^ from the extracellular matrix, (*ii*) as the astrocytic activity exceeds a critical threshold, K^+^ clearance from the extracellular matrix increases rapidly.

The coefficients *γ*_e_, *γ*_i_ represent the strengths of K^+^ release driven by optogenetically stimulated excitatory and inhibitory neurons, respectively. The coefficient *γ*_a_ represents the strength of K^+^ clearance driven by optogenetically stimulated astrocytes. These coefficients allow to encode population specific optogenetic effects in the model. They link the experimental manipulation via optogenetics to K^+^ dynamics.

See Table A1 for a summary of the model components and the corresponding control parameters.

**Table A1.**
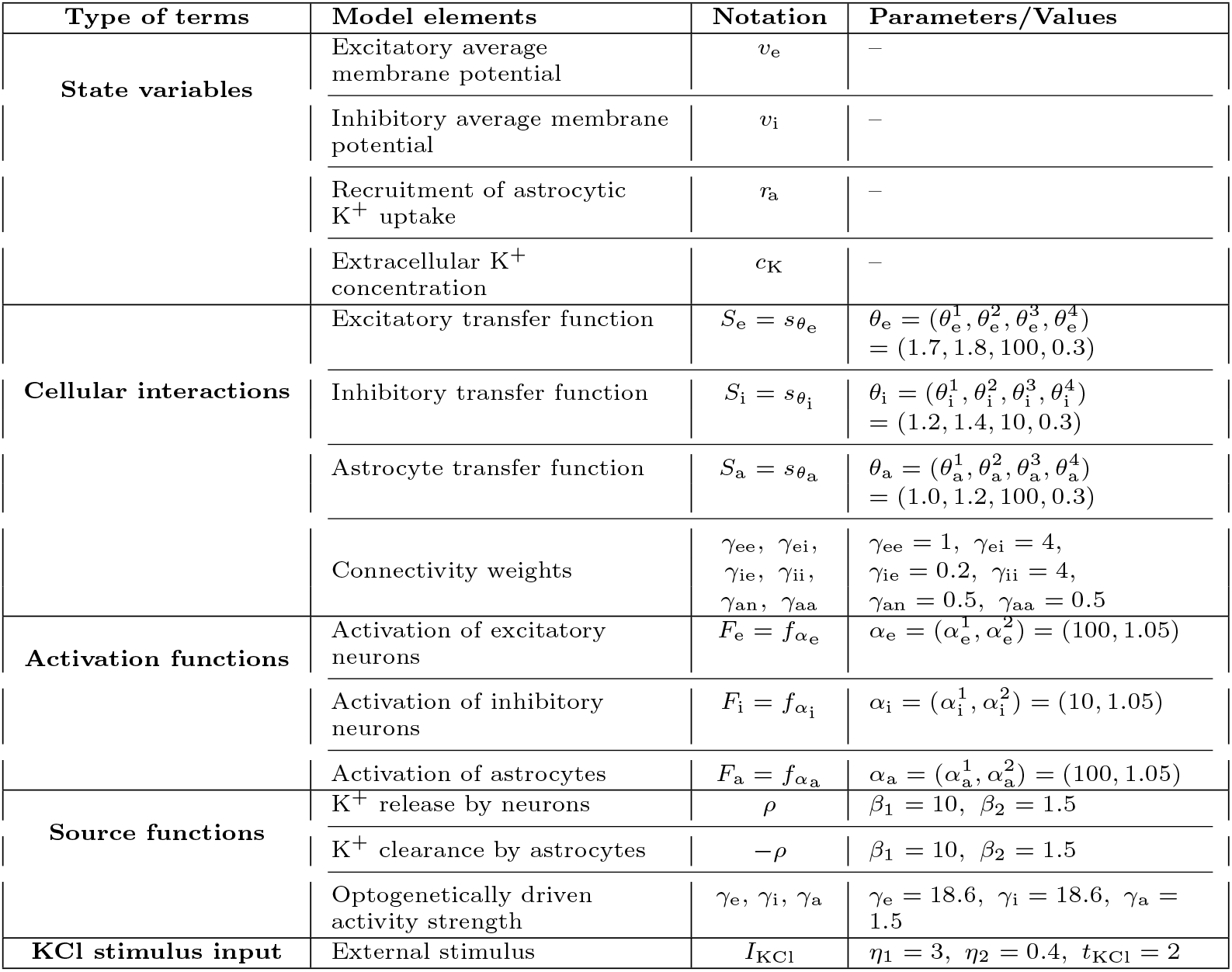
Model components. Right-hand entries indicate control parameter values.

## Appendix B

### Discretization

To numerically solve system (1), we discretize the cortical segment *D* using a regular spatial grid:

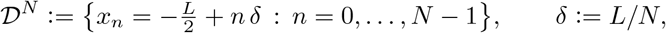

where periodic boundary conditions imply the identification *x*_0_ = *x*_*N*_. The Laplacian term *∂*^2^*c*_*K*_*/∂x*^2^ is approximated using the standard second-order finite-difference scheme.

The only subtlety lies in the efficient computation of the approximation of the convolution terms (3). After spatial discretization and at a given time, these terms reduce to:

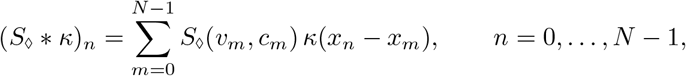

where *v*_*m*_ = *v*(*x*_*m*_) and *c*_*m*_ = *c*(*x*_*m*_). A direct evaluation of this sum has a computational cost of *O* (*N* ^2^) per time step, which becomes prohibitive for fine spatial grids. As the kernel *κ* is symmetric, it is possible to significantly reduce this cost, using the fact that, under periodic boundary conditions, the discrete convolution can be computed using the Discrete Fourier Transform (DFT). Introducing the vectors:

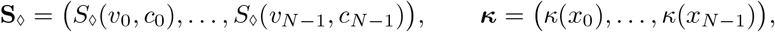

the discrete convolution is obtained via:

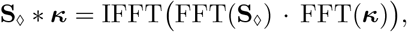

where FFT and IFFT denote, respectively, the Fast Fourier Transform and its inverse, and the product is componentwise.

The FFT-based approach reduces the computational complexity of the convolution from *O* (*N* ^2^) to *O* (*N* log *N*), providing substantial acceleration. To further optimize performance, the number of grid points *N* is chosen to be a power of 2, allowing the FFT algorithm to achieve its maximal efficiency.

This procedure yields an ordinary differential system of dimension 4 × *N*, which we integrate in time using the Matlab built-in solver ode45, an explicit fourth-order Runge–Kutta method with adaptive time-stepping.

## Appendix C

### CSD propagation speed in simulations

In the simulations, we study the CSD propagation by analyzing its propagation speed as the parameters corresponding to (EC1)-(EC4) are varied; see Appendix 3 for details regarding the experimental conditions (EC1)-(EC4). The propagation velocity *v* is computed in the simulations similarly to the experiments:

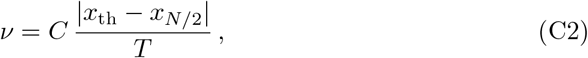

where *C >* 0 is a scaling constant, *T >* 0 denotes the final time of the simulation and *x*_*N/*2_ = 0 denotes the position at which a K^+^ puff is applied to trigger CSD. Here *x*_th_ is defined as

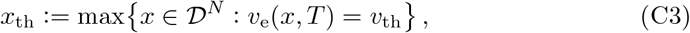

where *v*_th_ denotes a prefixed, sufficiently high potential threshold. Since CSD propagation is symmetric with respect to its initiation point *x*_*N/*2_ = 0 (Fig. 2), considering max or min in the definition of *x*_th_ does not change the results.

